# Guided Axon Outgrowth of Neurons by Real-Time Femtosecond Laser Penetration on a 4 μm Thick Thin-Glass Sheet

**DOI:** 10.1101/2021.08.17.456633

**Authors:** Dian Anggraini, Xun Liu, Kazunori Okano, Yo Tanaka, Naoyuki Inagaki, Ming Li, Yoichiroh Hosokawa, Sohei Yamada, Yaxiaer Yalikun

**Affiliations:** Division of Materials Science, Nara Institute of Science and Technology, Ikoma 630-0192, Japan; Center for Biosystems Dynamics Research (BDR), RIKEN, Osaka 565-0871, Japan; Division of Biological Science, Nara Institute of Science and Technology, Ikoma 630-0192, Japan; School of Engineering, Macquarie University, Sydney 2122, Australia

## Abstract

Transplantation of scaffold-embedded guided neurons has been reported to increase neuronal regeneration following brain injury. However, precise axonal integration between host and transplant neurons to form functional synapses remains a major problem. This study aims to develop a real-time femtosecond (fs) laser penetration on a 4 μm thick thin-glass sheet to promote guided axon outgrowth influenced by molecular gradients in a microfluidic device. The device enables the introduction of the guidance molecule (i.e., netrin-1), neuronal culture, and manipulation by fs laser. After fabricating multiple micro-holes on the thin-glass sheet using fs laser, netrin-1 gradients with radial concentrations are generated in the chamber, affecting axon outgrowth and guidance. A majority of axons (~92%) experiences guided outgrowth with positive angular changes towards netrin-1 gradients. These results demonstrate the capability of the precise and real-time manipulation system based on a fs laser and a microfluidic device to control the growth of neurons.

## Introduction

In the last decades, the high incidence of brain injury cases and the low rates of successful recovery have become serious health problems (Corrigan et al., 2010; Dohmen et al., 2008; Lee et al., 2016; Ma et al., 2014; Pedersen et al., 1995). Cell transplantation is the current gold standard for treating brain injuries, especially those identified by large lesions, such as traumatic brain injury, spinal cord injury, and stroke (Fan et al., 2010; Hawryluk et al., 2014; Kondziolka et al., 2000; Pearse et al., 2004; Riess et al., 2002; Shear et al., 2004; Shin et al., 2015). However, after the transplantation of cells alone, neuronal regeneration remains one of the greatest challenges, as identified by poor cell survival, differentiation, and axonal integration (Fan et al., 2010; Hawryluk et al., 2014; Shear et al., 2004; Vroemen et al., 2003). Precise axonal integration to form functional synapses is the ultimate goal to obtain functional recovery after brain injury (Hollis et al., 2015; Yokota et al., 2015). Thus, it requires the incorporation of tissue engineering approaches to promote functional improvements following brain injury. Transplantation of scaffold-embedded guided axon outgrowth has been reported, which can increase cell survival, axon outgrowth, neuronal tissue reconstitution, and decrease inflammation (Carlson et al., 2016; Vaysse et al., 2015; Zhang et al., 2018). Transplanted scaffold can also be manipulated to release supporting factors, such as growth factor and trophic factor, to increase regeneration rate following brain injury (Kadoya et al., 2016; Lu et al., 2012). Apart from its functionality, scaffold that fulfills thin and biocompatible characteristics is also important to prevent collateral damages among host cells and tissues during transplantation (Nayak et al., 2010; Prabhakaran et al., 2008; Tao et al., 2007; Tao and Desai, 2007). While many transplanted scaffolds with different materials have been developed (Carlson et al., 2016; Vaysse et al., 2015; Zhang et al., 2018), glass is the most favorable material to fulfill the need.

Recently, advanced methods, such as topographic and chemical surface patterning, have been developed to regulate axon outgrowth on the scaffold (Alsmadi et al., 2015; Dinis et al., 2014; Kador et al., 2014; Millet et al., 2010; Yang et al., 2017). The manipulation to assemble guided neurons on the scaffold is performed in vitro, followed by transplanting a scaffold containing fully guided neuronal networks into the lesion in brain injury (Carlson et al., 2016; Vaysse et al., 2015; Zhang et al., 2018). These approaches increase the efficacy of neuronal regeneration, however, precise axonal integration between transplant and host neurons to form functional synapses remains a formidable challenge. Therefore, it calls for a high-precision method for direct axon outgrowth in real-time conditions to attain robust axon regeneration and promote functional recovery after brain injury.

Femtosecond (fs) laser is a powerful tool for precisely manipulating cells and scaffolds (Hosokawa et al., 2009; Kaji et al., 2007; Okano et al., 2016, 2013, 2011; Rukmana et al., 2019; Yamamoto et al., 2011). Fs laser can be positioned in sophisticated optical arrangements that allow the placement of other controllers (e.g., a motorized stage and an environmental chamber), resulting in reliable and reproducible experiments. Also, fs laser offers spatiotemporal flexibility to control the objects in different sites and timing, realizing manipulation in real-time conditions and at single-cell resolutions (Hosokawa et al., 2011, 2009; Rukmana et al., 2019). Compared to other lasers, fs laser processing produces low heat which can minimize thermal-induced cell death (Kasaai et al., 2003; Le Harzic et al., 2002). Based on these favorable characteristics, our group has successfully developed guided neurite outgrowth by ablating the cytophobic area to expose the cytophilic area of the scaffold, resulting in neurites grow precisely along with the linear shape of the ablated area (Okano et al., 2016, 2013; Yamamoto et al., 2011). However, these studies have limitations for transplantation purposes in donating molecules to control axon outgrowth of neurons in real-time conditions.

In this study, we developed a precise and real-time fs laser penetration on a 4 μm thick thin-glass sheet to allow guided axon outgrowth by molecular gradients in a microfluidic device. We used a thin-glass sheet (4 μm thickness) as a scaffold (Yuan et al., 2021), which has been used for precise sensing and controlling fluids in microfluidic devices (Tang et al., 2021; Yalikun and Tanaka, 2017). More importantly, penetrated micro-holes or channels can be easily and precisely fabricated using fs laser on a thin-glass sheet (Yalikun et al., 2017, 2016). To the best of our knowledge, this study is the first to control axon outgrowth of primary culture of hippocampal neurons based on laser-fabricated micro-holes on a thin-glass sheet (4 μm thickness). Hippocampal neurons isolated from embryonic day 16 (E16) mice were cultured on the thin-glass sheet for 1 day in vitro (DIV), while the axon guidance molecule (i.e., netrin-1) (Ishii et al., 1992; Serafini et al., 1994) was manually introduced to the channel. The areas of a thin-glass sheet near the neurons that have axon projections were selected to focus fs laser pulses to create the micro-holes. Netrin-1 gradients released from multiple micro-holes affected the growth and guidance of the cultured neurons. A majority of axons (~92%) were guided toward the netrin-1 gradients, indicated by narrower axonal angular changes during exposure to netrin-1. The guidance process occurred at different guided times, resulting in different axon elongations depending on the distance of axons to the micro-holes. In addition, the dynamics of guided axon outgrowth were observed starting from 1 to 16 h exposure to netrin-1. These results demonstrated that the developed system is capable of controlling guided axon outgrowth precisely in real-time conditions (Figure 1). We believe that our system can be used to study real-time and detailed mechanisms of development and regeneration of nervous systems. Further, this system can be applied to precisely regulate guided axon outgrowth in the actual lesion in brain injury.

**Figure 1.**
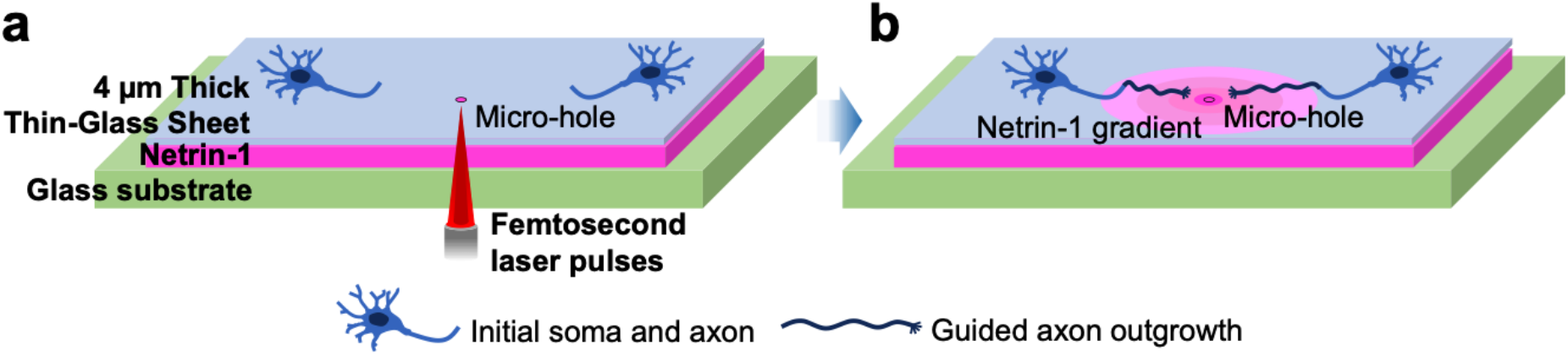
Conceptual illustration of real-time femtosecond (fs) laser penetration on a 4 μm thick thin-glass sheet for guiding axon outgrowth in a microfluidic device. (a) Pulses of fs laser irradiate the thin-glass sheet to fabricate the micro-hole during neuronal culture. (b) A netrin-1 gradient generated from the laser-fabricated micro-hole guides axon outgrowth of hippocampal neurons towards the micro-hole.

## Results

### Design and principle of the microfluidic device

The microfluidic device enables the embedding of culture substrate (i.e., 4 μm thick thin-glass sheet) penetrated by fs laser pulses to generate a simple diffusion process of a guidance molecule from the channel to the chamber. The microfluidic device was comprised of five layers from the bottom to the top, i.e., a glass substrate, a channel, a thin-glass sheet, 3 chambers, and 2 PDMS slabs. The width of the channel and chamber was 3 mm and 1 mm, respectively, allowing the thin-glass sheet (4 mm width) to separate the compartment for cell culture and guidance molecule. The chamber with a width of 1 mm was fabricated to confine the growth area of the cells on a thin-glass sheet (Figure 2a). This limited growth area was convenient for observing cell behaviors in a single field of view microscope. The chamber and PDMS slabs depth were sufficiently high (2 mm in total) to ensure the established long-term molecular gradient in the chamber (Figure 2a and b). It was also favorable to retain the amount of the medium in a microfluidic device. For long-term cell culture, a five-layered device was placed inside a dish containing medium and a wet paper (Figure 2c). This setup can maintain cell culture in a stable pH medium without evaporation for up to 7 DIV.

**Figure 2.**
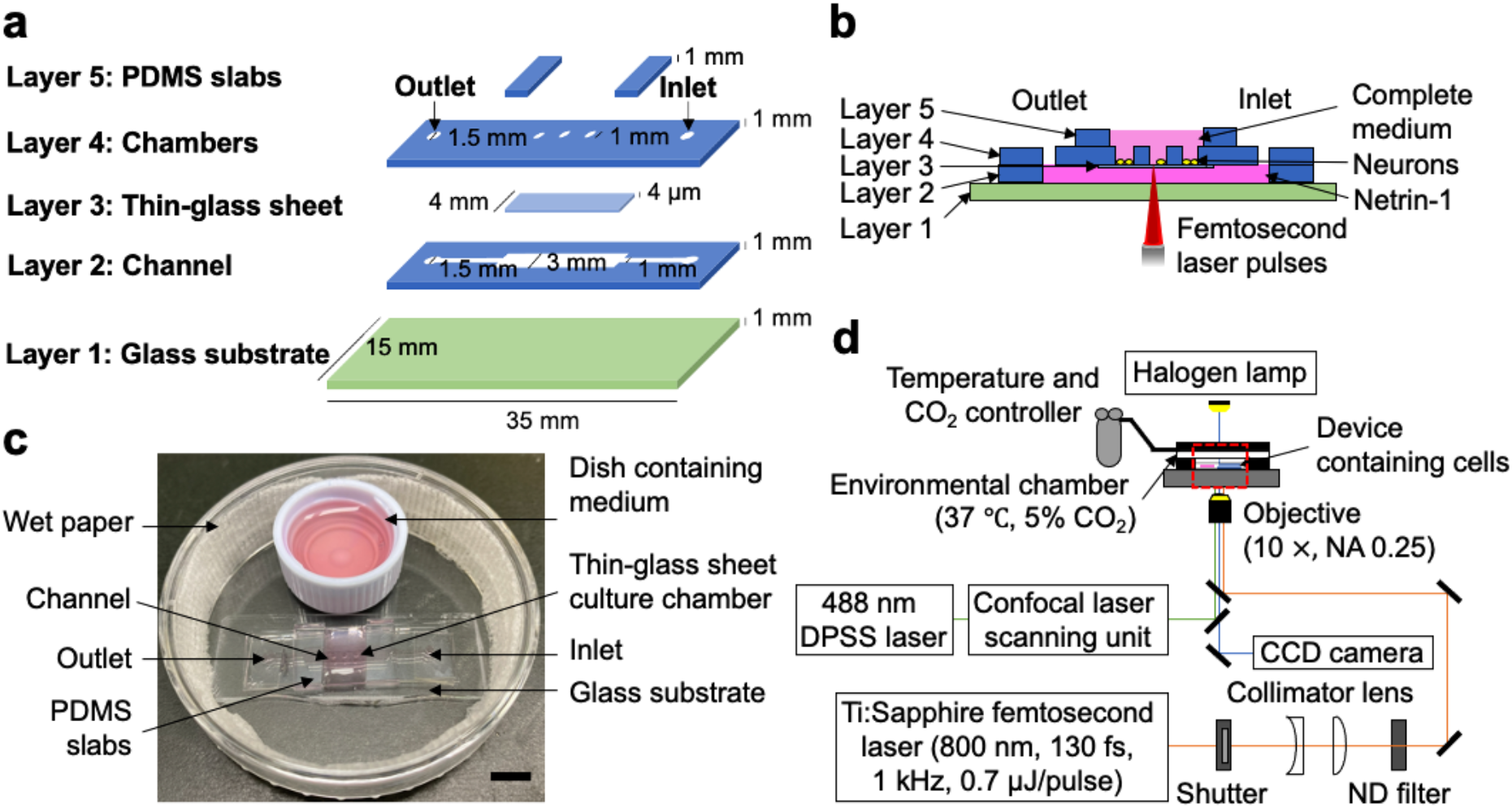
Design and principle of the microfluidic device and femtosecond (fs) laser penetration on a 4 μm thick thin-glass sheet embedded in a microfluidic device. (a) Schematic illustration of a five-layered microfluidic device for real-time neuronal manipulation using fs laser to control axon outgrowth. (b) A side view of the microfluidic device containing neurons, guidance molecule (i.e., netrin-1), and complete medium. Pulses of fs laser irradiate a thin-glass sheet (4 μm thickness) to fabricate micro-holes to release netrin-1, inducing the response of axon guidance. (c) The microfluidic device is placed inside a dish containing medium and a wet paper to prevent medium evaporation. (d) Schematic illustration of real-time fs laser processing and time-lapse imaging. The scale bar is 5 mm.

### Characterization of molecular gradient

To characterize molecular gradient in the microfluidic device, we first performed the finite element method simulation to model the diffusion pattern of guidance molecules released from laser-fabricated micro-holes over time. The guidance molecule diffused from the channel to the chamber with multiple radial concentrations via the micro-holes (Figure 3a). The guidance molecule was continuously released from the micro-holes over time (Figure 3a and Figure supplement 2). In parallel, we also conducted the experimental characterization of a molecular gradient with Alexa Fluor 488-BSA to confirm concentration profiling of a guidance molecule over time. Molecular weight of BSA (66.0 kDa) was similar to netrin-1 (68.2 kDa) which was expected to have a similar molecular gradient pattern. The surface of the thin-glass sheet was selected as the focused area to observe the fluorescence intensity changes, representing the location of the axon outgrowth responses to the molecular gradient (Figure 3b). We introduced the Alexa Fluor 488-BSA to the channel located on the below side of the focused area, defining reference fluorescence for measurements. The fluorescence intensity was continuously taken from 1 to 16 h after laser processing (Figure 3c). Fabrication of 9 micro-holes with fs laser pulses on the thin-glass sheet was carried out for 1 h, so that the fluorescence observation after laser processing was started after the micro-holes are all fabricated. The fluorescence intensity increased by 9 h and attained a plateau after 10 h (Slope = 0.32 μm^-1^, R^2^ = 0.73, *p* = 1.85 × 10^-29^, *p* < 0.05, Regression) (Figure 3d). Moreover, the molecule diffused and generated a gradient from the area near (0 μm) to far from the micro-hole (450 μm) over time (Figure 3e). The fluorescence intensity shift was higher at the early time-point (1 h) than at the final time-point (16 h) (Figure 3f). The series of results show the capability of the microfluidic device with fs laser pulses for generating molecular gradients to induce guided axon outgrowth.

**Figure 3.**
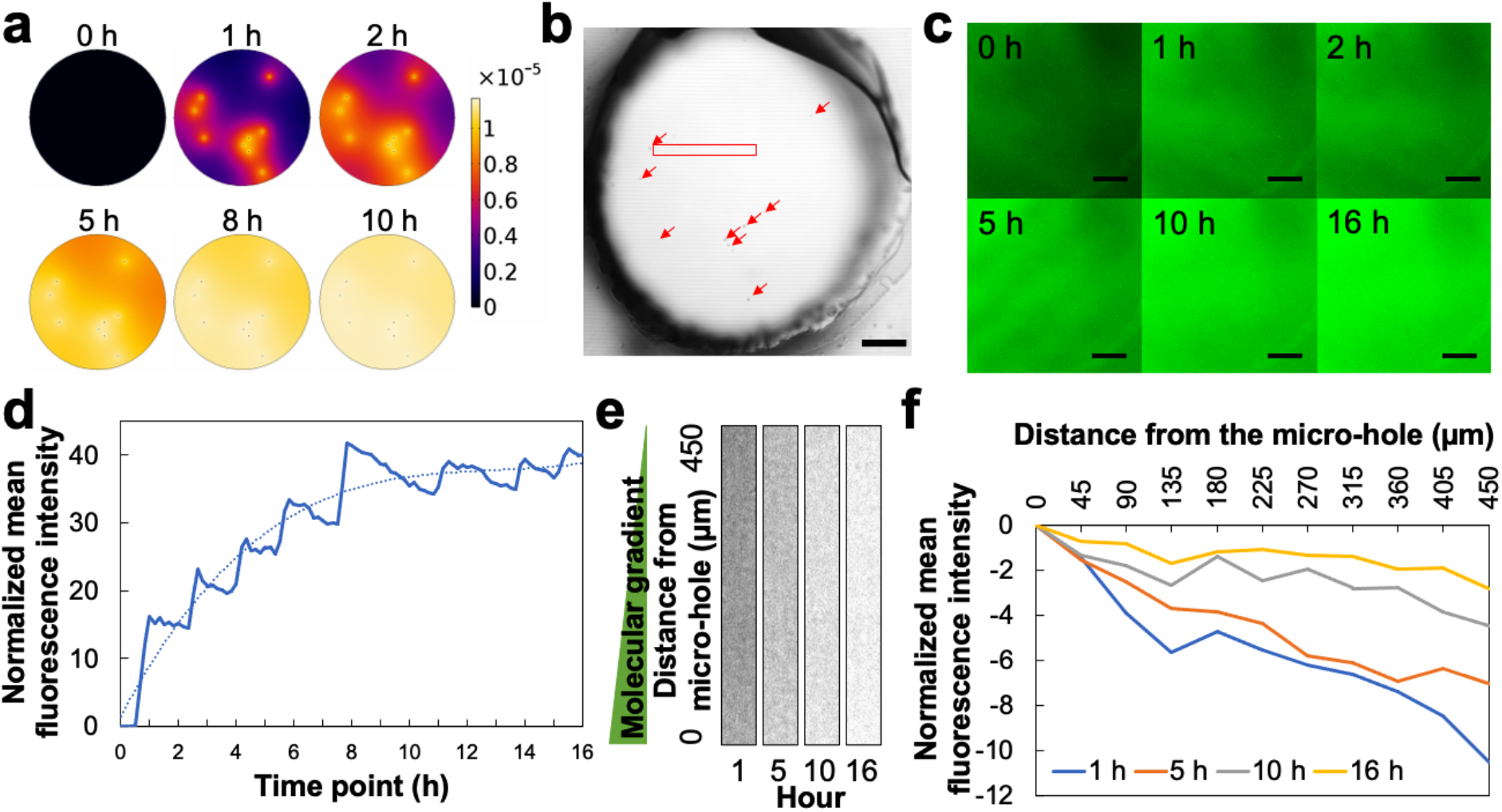
Simulation and experimental characterization of molecular gradients. (a) Twodimensional (2D) diffusion simulation of molecular gradients in the chamber over time. (b) An image of the thin-glass sheet surface covered by the chamber as the focused area to observe fluorescence intensity. Red arrows indicate the position of laser-fabricated micro-holes. (c) Fluorescence images of molecular distribution in the chamber over time. The fluorescence images are observed at a single-layer before (0 h) and after the laser processing (1 h, 2 h, 5 h, 10 h, and 16 h). (d) Fluorescence intensity shift in the chamber for 16 h. (e) Fluorescence images of a molecular gradient from a red rectangle in (b) over time. (f) Fluorescence intensity shift of a molecular gradient across the field at 1 h, 5 h, 10 h, and 16 h. The scale bars are 200 μm in (b) and (c).

### Real-time manipulation system by femtosecond laser pulses

To the best of our knowledge, this study is the first to utilize a thin-glass sheet (4 μm thickness) as a culture substrate for hippocampal neurons. The thin-glass sheet (4 μm thickness) has recognized as the thinnest glass with high transparency (Figure 4a), which is beneficial for use as a culture substrate manipulated by laser technology. The extracellular matrix (i.e., laminin) was coated on the thin-glass sheet surface for a cell scaffold. Neurons adhered to the thin-glass sheet surface at 3 h after cell seeding, then started to project their axons at 1 DIV. This state of growth was suitable for manipulation by fs laser to prevent cell detachment.

**Figure 4.**
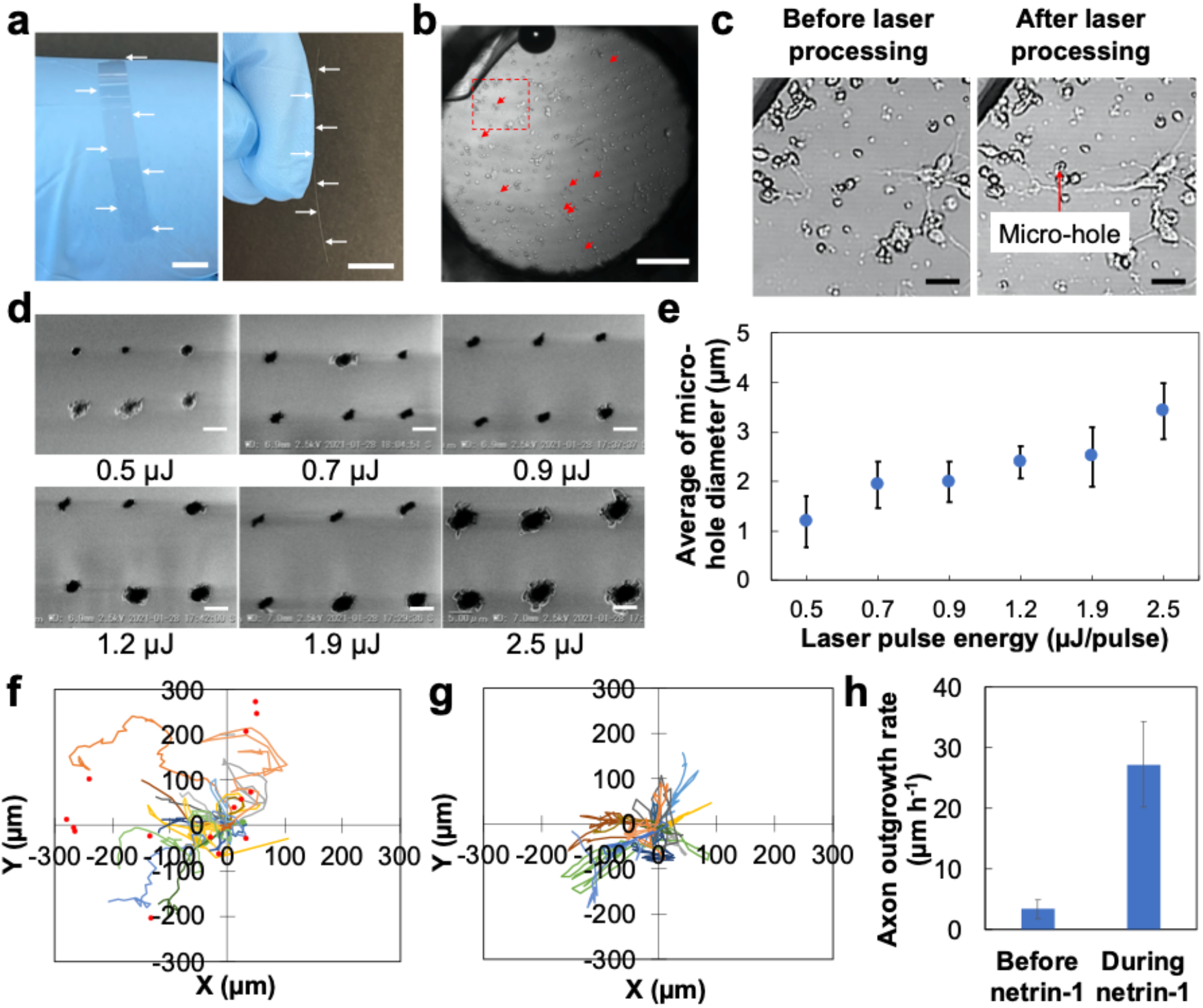
Scanning Electron Microscopy (SEM) observation of laser-fabricated micro-holes on a 4 μm thick thin-glass sheet and response of the hippocampal neurons to the netrin-1 gradients. (a) A top (left) and side view of the thin-glass sheet (right) showing its ultra-thin thickness. (b) A photomicrograph of the 1 DIV neuronal culture. Red arrows indicate the laser-fabricated micro-holes. (c) Photomicrographs from the red dashed line in (b) before (left) and after laser processing (right) (d) A top view of SEM images of laser-fabricated micro-holes on the thin-glass sheet by different pulse energies: 0.5, 0.7, 0.9, 1.2, 1.9, and 2.5 μJ/pulse. (e) Laser pulse energy dependency on the laser fabricated micro-hole diameter of the thin-glass sheet obtained by different pulse energies. (f and g) Axon trajectories of neuronal culture with netrin-1 gradients generated from laser-fabricated micro-holes (f) and without netrin-1 for 16 h (g). The initial axons are normalized at (x = 0 μm and y = 0 μm). Red dots are the normalized positions of the micro-holes. (h) Axon outgrowth rate before and during exposure to netrin-1. Values are shown as means ± SEM (n = 20). The scale bars are 5 mm in (a), 200 μm in (b), 50 μm in (c), and 5 μm in (d).

The areas of the thin-glass sheet near the projected axons were selected as the laser-focused spot (Figure 4b). Various laser pulses were examined to obtain a short-time laser processing and minimal water bubble generation to prevent cell interferences. The laser pulse of 0.7 μJ/pulse was successful in fabricating the micro-holes with a short-time laser penetration (i.e., 2 – 8 s/micro-hole) and minimal water bubble generation (Video supplement 1). As a result, nine micro-holes were fabricated on the thin-glass sheet near the projected axons. The fabrication conditions with 0.7 μJ/pulse fs laser did not affect cell detachment and damage (Figure 4c).

Scanning Electron Microscopy (SEM) images confirmed the diameter of the microholes fabricated using fs laser pulses of 0.7 μJ/pulse ranges from 1.11 to 2.75 μm. Furthermore, SEM observations were also carried out to investigate the diameter of the micro-holes fabricated with different laser pulse energies (Figure 4d and Figure supplement 3). Micro-hole diameter averages were: 1.19 ± 0.51, 1.93 ± 0.40, 1.98 ± 0.31, 2.39 ± 0.60, 2.50 ± 0.56, and 3.42 ± 0.51 μm, when the laser pulse energies were 0.5, 0.7, 0.9, 1.2, 1.9, and 2.5 μJ/pulse, respectively (Figure 4d and e). The SEM analysis reveals that micro-hole diameter increased with the increase of the laser pulse energy.

### Response of the hippocampal neurons to the netrin-1

After demonstrating the capability of the device for generating molecular gradients, we utilized this device for regulating axon outgrowth by an axon guidance molecule (i.e., netrin-1). Through the micro-holes fabricated by fs laser pulses, netrin-1 was released and distributed throughout the chamber over time, influencing neuronal responses. Axons with varying distances to the netrin-1 source (micro-holes) were investigated before (0 h) and after laser processing (1 – 16 h). In cultures exposed to netrin-1 for 16 h, individual axon trajectories showed a tendency for axons to grow toward the micro-holes, while no directional trend was shown in cultures without netrin-1 (Figure 4f and g). The axons experienced outgrowth during exposure to netrin-1, with a mean axon outgrowth rate of 27 μm/h (Figure 4h).

### Angular changes of netrin-1-responsive neurons

To further determine whether the neurons responded to the netrin-1 source, angular changes of axons before and after laser processing were evaluated. Angular changes were analyzed by subtracting angles of axonal end-points toward the micro-holes before and during exposure to netrin-1 (Figure 5a and Figure supplement 1). The observed axons had varying angles, identified as axons at angles of ≤ 30°, ≤ 60°, ≤ 90°, ≤ 120°, ≤ 150° toward the micro-holes before exposure to netrin-1. After the laser processing, guided axon outgrowth to the netrin-1 source was investigated at different time points depending on the farthest axon elongation toward the micro-holes, resulting in 2 axons at ≤ 30° persisted at ≤ 30° toward the micro-holes; 7 axons at ≤ 60°, 2 axons at ≤ 90°, 3 axons at ≤ 120°, and 1 axon at ≤ 150° turned into ≤ 30° toward the micro-holes; and 2 new axons grew with an angle of ≤ 30° toward the micro-holes (Figure 5b, d, and e). The remaining axons were 5 axons at ≤ 60° and 2 axons at ≤ 90° toward the micro-holes (Figure 5b). The mean angle of axon turning during exposure to netrin-1 was 41°. Based on this average, 22 out of 24 axons turned with positive angular changes toward the micro-holes: 11 axons turned more than 41° toward the micro-holes, 11 axons turned less than 41° toward the micro-holes, and the remaining 2 axons turned away from the micro-holes with negative angular changes (*p* < 0.0001, Independent t-test) (Figure 5c). Therefore, ~92% of axons showed guided toward the micro-holes indicated by positive axonal angular changes. Apart from the phenomenon of guided axon outgrowth, some guided axons also crossed with other guided axons and preexisting axons in the areas surrounding the micro-holes (Figure supplement 4). These results showed that the axons respond to the netrin-1 sources, and demonstrated the reliability of the system for regulating the process of guided axon outgrowth following the fabrication of micro-holes by fs laser.

**Figure 5.**
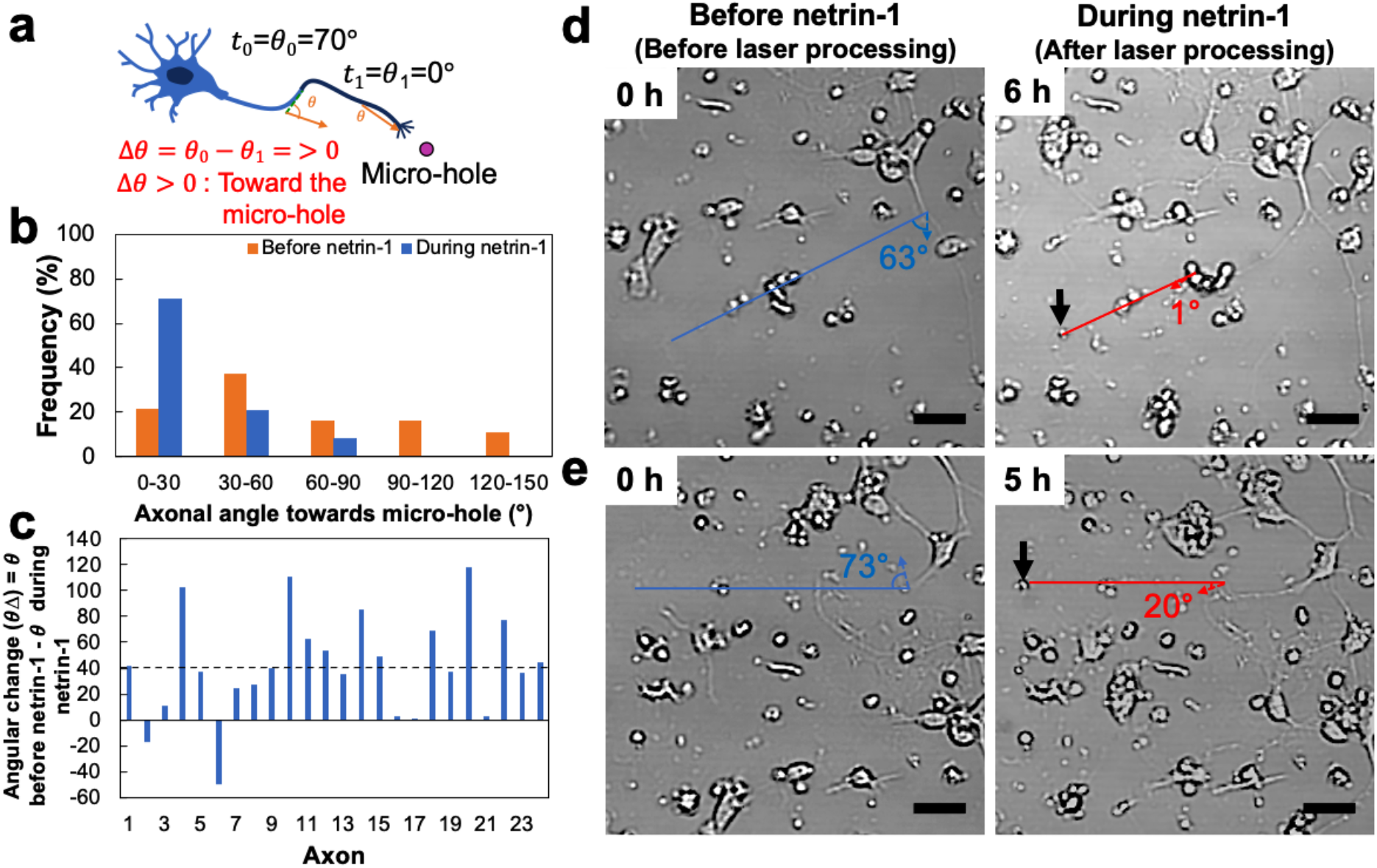
Angular changes of netrin-1-responsive neurons. (a) Schematic illustration of the angular change (Δ*θ*) of the initial axon angle (*θ*_0_) before micro-hole fabrication (t_0_) with axon outgrowth angle (*θ*_1_) after micro-hole fabrication (t_1_). (b) Frequency of axonal angle distributions towards the micro-holes before and during exposure to netrin-1. (c) Angular change of axons before and during exposure to netrin-1. Dashed line represents the mean angle of turning (41°) (*p* < 0.0001, Independent t-test, n = 24). (d and e) Photomicrographs of axonal angle towards the micro-hole before (left) and during exposure to netrin-1 (right), showing narrower axonal angular changes. Black arrows are the micro-holes. The scale bars are 50 μm.

### Growth dynamics of netrin-1-responsive neurons

As noted above, netrin-1 released from laser-fabricated micro-holes can induce growth and guidance of axons of hippocampal neurons. We next investigated the growth dynamics of netrin-responsive neurons before and after exposure to netrin-1. The axons were categorized based on their angles to the micro-holes, i.e., toward micro-holes (axons at angles of ≤ 45°) and away from micro-holes (axons at > 45°). The axons grew with the rate ranging from 0 to 190 μm/h (Figure 6a). The highest incidence of axon outgrowth rate was 0 – 10 μm/h. Based this outgrowth rate, the axons grew away from the position of micro-holes before exposure to netrin-1. After the micro-holes were fabricated, the outgrowth of guided axons toward the micro-holes continued to increase from 1 to 16 h netrin-1 exposure (Figure 6b). These results indicate that netrin-1 released via the micro-holes induces the dynamics of guided axon outgrowth for 16 h.

**Figure 6.**
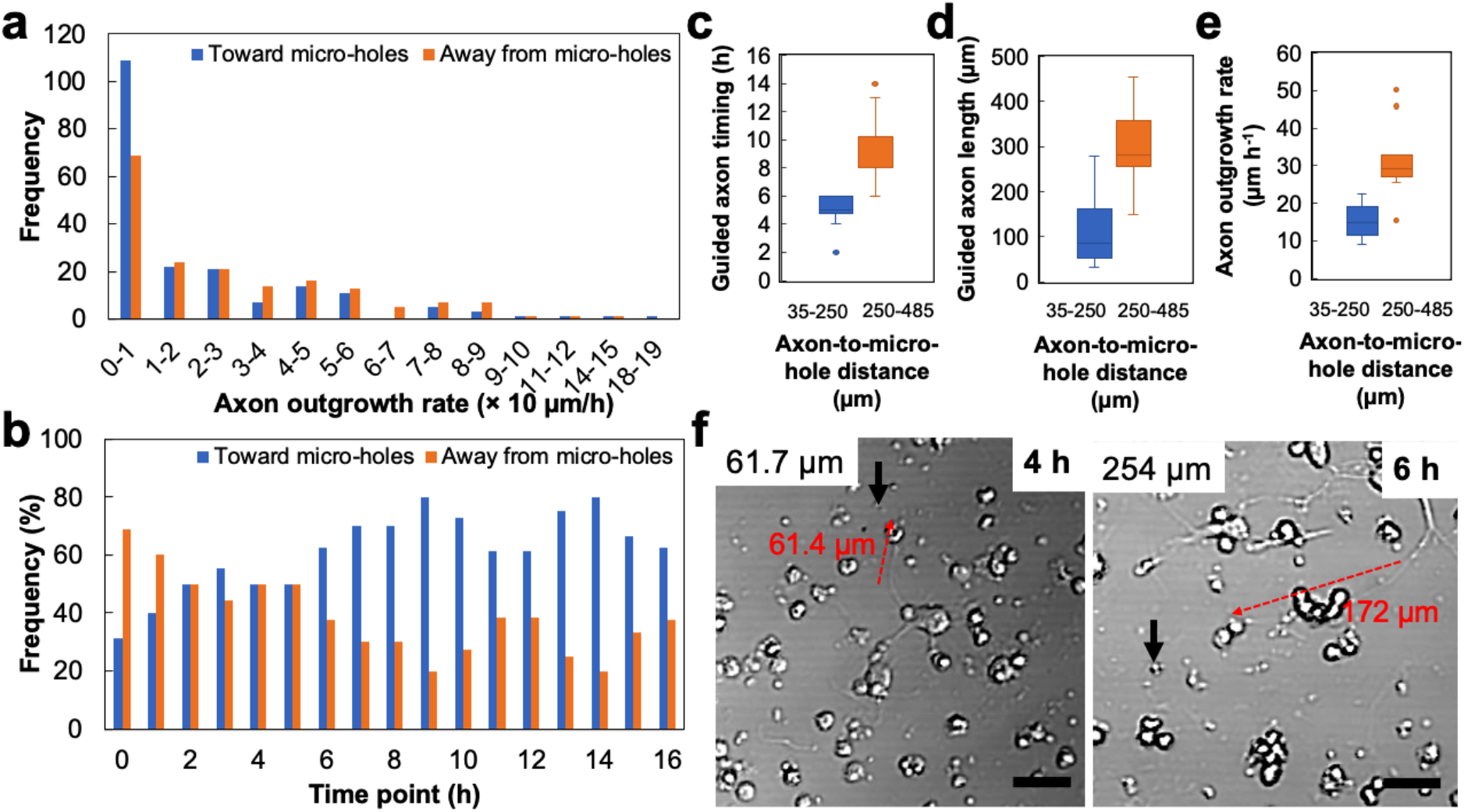
Growth dynamics of netrin-1-responsive neurons and distance-dependent neuronal response to netrin-1. (a) Frequency of axons grows toward the micro-holes (blue) and away from the micro-holes (orange) at different axon outgrowth rates. (b) Frequency of axons grows toward the micro-holes (blue) and away from the micro-holes (orange) at an axon outgrowth rate of 0 – 10 μm/h. (c, d, and e) Dependency of axon-to-micro-hole distance against guided axon timing (c), guided axon length (d), and axon outgrowth rate (e) located near (35-250 μm) and far from the micro-holes (250-485 μm) (*p* < 0.001, Independent t-test, n = 24). (f) Photomicrographs of axon outgrowth to the micro-holes (red dashed arrow line) for axons located near (61.7 μm) (left) and far from the micro-holes (254 μm) (right). Black arrows are the micro-holes. The scale bars are 50 μm.

### Distance-dependent neuronal response to netrin-1

We finally analyzed the axons’ response dependent on the distances toward the molecular sources. The initial axons with varying distances to the micro-holes were evaluated based on guided time and axon length. Guided time was the time needed for axons to grow farthest toward the micro-holes. The initial axons elongated toward the molecular gradients with varying lengths at different time points during exposure to netrin-1. The graphics depicted that the guided time and axon length for the axons located closer to the micro-holes (35-250 μm) were shorter than those of the axons located far from the micro-holes (250-485 μm) (*p* < 0.0001; *p* < 0.001, Independent t-test, respectively) (Figure 6c, d, f, Video supplement 2, and Video supplement 3). Interestingly, axons located near the micro-hole also showed a lower axon outgrowth rate than those far from the micro-holes (Figure 6e). These analyses demonstrate that guided time, guided axon length, and axon outgrowth rate are dependent on the distance of axons to the micro-holes.

## Discussion

Here, we demonstrated a precise and real-time neuronal manipulation for controlling axon outgrowth enabled by guidance molecule released from fs laser-fabricated micro-holes on a 4 μm thick thin-glass sheet. The real-time manipulation was achieved by penetrating the thin-glass sheet using fs laser during cell culture in a microfluidic device. The microfluidic device was designed without geometric limitations (e.g., microchannel arrays and asymmetric channels) (Gladkov et al., 2017; Vaysse et al., 2015; Yang et al., 2017), allowing free and random cell growth on the culture substrate. Thus, the device relied solely on generated radial concentrations of molecular gradients by laser-fabricated micro-holes, inducing neuronal responses toward the gradients (Figure 3, 4f, 4h, and 5). Radial concentrations were identified as a typical pattern for local manipulation systems, such as micropipette-based chemotactic assay and implant-based drug delivery (Homsy et al., 2015; Shi et al., 2019). Unlike other microfluidic devices that utilize fluidic controllers (e.g., pumps and valves) to generate fluid flow (Bhattacharjee and Folch, 2017; Chen et al., 2012; Millet et al., 2010; Nam et al., 2007; Xiao et al., 2013), this device enabled the diffusion of guidance molecule from the channel to the chamber via laser-fabricated micro-holes without any concerns of fluid flow-induced cell interferences.

To the best of our knowledge, this study is the first to show the response of hippocampal neurons to netrin-1 released from fs laser-fabricated micro-holes. The gradient of netrin-1 with concentrations ranging from 5 to 1000 ng/mL has been reported to induce axon outgrowth of primary neurons (Baba et al., 2018; Bhattacharjee et al., 2010; Bhattacharjee and Folch, 2017; Blasiak et al., 2015; Kothapalli et al., 2011; Serafini et al., 1994; Taylor et al., 2015; Xiao et al., 2013; Xu et al., 2018). In this current study, netrin-1 (800 ng/mL) was introduced to the channel. The netrin-1 in the channel diffused via micro-holes to the complete medium in the chamber, resulting in an accelerated axon outgrowth rate of neurons during exposure to netrin-1 for 16 h (Figure 4h). The mean axon outgrowth rate (27 μm/h) was comparable to other experiments with a threefold extension from their soma (Bhattacharjee et al., 2010; Bhattacharjee and Folch, 2017; Kothapalli et al., 2011).

Apart from the role of netrin-1 to promote the outgrowth rate of neurons, netrin-1 has been known to organize guided axon outgrowth on various scaffolds in soluble and scaffold-attached form (Baba et al., 2018; Bhattacharjee and Folch, 2017; Kador et al., 2014; Moore et al., 2012, 2009; Xiao et al., 2013). In soluble form, netrin-1 is introduced in a one-sided manifold to control axon outgrowth (Baba et al., 2018; Bhattacharjee and Folch, 2017; Xiao et al., 2013). In scaffold-attached form, netrin-1 is immobilized in a scaffold to form a topographic gradient to guide the outgrowth of axons (Kador et al., 2014; Moore et al., 2009). Unlike those systems, this study achieved local manipulation activated by multiple micro-holes-generated netrin-1 gradients to induce precise guided axon outgrowth (Figure 3), where narrower axonal angular changes toward the micro-holes were observed (Figure 5b, d, and e). A large majority of the axons (~92%) experienced positive angular changes during exposure to netrin-1 gradients at specific time-point (Figure 5c). Moreover, the dynamics of guided axon outgrowth for 16 h were shown in the trajectory of axons directed toward the micro-holes (Figure 4f and 6b). The long-term guidance effect of netrin-1 was in accordance with the release of netrin-1 from 0 to 9 h and attain a plateau after 10 h (Figure 3d and Figure supplement 2).

Apart from being guided toward the micro-holes, some axons crossed with other guided axons and preexisting axons (Figure supplement 4). This phenomenon has been confirmed as synaptic connections in another local manipulation for targeting neuronal connections to form neuronal networks (Oyama et al., 2015; Taylor et al., 2010). Thus, physical and chemical cue are needed to direct the axons to connect with specific targets (Braisted et al., 2000; Nagel et al., 2015; Oyama et al., 2015). Furthermore, guided axon outgrowth was observed at initial axons located near and far from the micro-holes-generated-netrin-1 gradients. The initial axons near the micro-holes received the netrin-1 gradients earlier with higher concentrations than those far from the micro-holes, affecting guided axon outgrowth with a shorter guided time. In addition, the axons also had a shorter guided length with an average outgrowth rate lower than the axons far from the micro-holes (Figure 6c, d, e, f, Video supplement 2, and Video supplement 3). These shorter responses are caused by the saturation effect, where receptors in the axonal growth cones respond to high concentrations of netrin-1 (Keino-Masu et al., 1996; Ming et al., 1997).

In sum, the combination of the femtosecond laser and microfluidic device approaches can achieve guided axon outgrowth with precise, non-invasive, and real-time manipulation. The femtosecond laser setup facilitates the controllable system to fabricate micro-holes, generating the slow stream of molecules to induce the growth of slow-response cells, especially neurons. The microfluidic device enables the storing of guidance molecule that is important for real-time manipulation of cellular phenomena, i.e., guided axon outgrowth. Moreover, the use of passive microfluidics prevents shear stress-induced cell interferences, allowing to obtain exact cell information. The 4 μm thick thin glass sheet embedded in a microfluidic device enables the simple process of extracellular matrix coating, which acts as a culture substrate for neurons. The ultra-thin thickness of the glass allows short-time manipulation by laser technology, providing non-invasive cell manipulation. Also, the opacity property of the thin-glass sheet improves the accuracy for laser manipulation and increases the resolution for observing adherent cells. Based on these favorable characteristics, we believe that this system can be applied to investigate other cell types for different experimental purposes, e.g., neuron regeneration, immune cell response, stem cell differentiation, cancer cell proliferation, wound healing process, and other biological phenomena. For future applications, this system is expected to be developed as on-field neuronal transplantation to obtain robust axon regeneration and promote functional recovery in brain injury cases.

## Materials and Methods

### Fabrication of microfluidic device

A microfluidic device in this system allowed the introduction of guidance molecules and real-time manipulation using fs laser. The microfluidic device consisted of five layers from the bottom to the top. Layer 1 was a glass substrate (1 mm thickness) used as the basis of the microfluidic device. Layer 2 was a channel (1 mm depth and 3 mm width) for placing the guidance molecule (i.e., netrin-1). Layer 3 was a thin-glass sheet (4 μm thickness and 4 mm width) used as a culture substrate and was penetrated by fs laser pulses to fabricate the microholes. Layer 4 were three chambers (1 mm depth and 1 mm diameter) for confining the growth area of the cells and placing the medium. Layer 5 were two polydimethylsiloxane (PDMS) slabs (1 mm thickness) placed opposite the chambers to prevent the outspread of the medium. Inlet and outlet were fabricated directly to connect with the channel (Figure 2a). The thin-glass sheet was fabricated by weight-controlled load-assisted precise thermal stretching (Yuan et al., 2021). The chamber, channel, and PDMS slabs were fabricated using standard soft lithography fabrication. All the compartments were bonded together by plasma treatment (Plasma Cleaner CY-P2L-B) to form a five-layered device (Figure 2b). The device was then dry sterilized (As ONE SONW-300) at 160 °C for 1 h to prevent contamination.

### Simulation of gradient generation

Simulation of netrin-1 diffusion from the channel to the chamber via laser-fabricated micro-holes was performed using the commercial finite element software COMSOL Multiphysics 5.4 (http://www.comsol.com). The simulations were conducted using both two-dimensional (2D) and three-dimensional (3D) models. In the chamber, there was no convection; thus, the diffusion of netrin-1 from the channel to the chamber was described by diffusion equations:

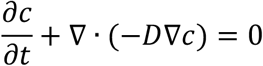

where t denoted time, c was the concentration, and D was the solute diffusion coefficient, and D in an aqueous culture medium was 2.36 ×10^-12^ m^2^/s. Concentration boundary condition occurred in the channel, defined as c=c0, c0=1×10^-5^ mol/m^3^. These were defined as no flux boundary conditions:

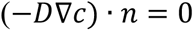

### Characterization of gradient generation

The diffusion of netrin-1 from the channel to the chamber via laser-fabricated micro-holes was confirmed with bovine serum albumin (BSA) labeled Alexa Fluor 488 (Invitrogen, USA). Alexa Fluor 488-BSA (1.77 μM) was introduced to the channel of the device. The distribution of Alexa Fluor 488-BSA was observed with time-lapse imaging using a confocal fluorescence microscope. A 488 nm diode-pumped solid-state (DPSS) laser (Spectra-Physics, PC14763) was used to excite Alexa Fluor 488-BSA. The observation was conducted before (0 h) and after laser processing (1 – 16 h). The fluorescence intensity shift of the Alexa Fluor 488-BSA in the chamber was evaluated by subtracting the fluorescence intensity value after laser processing (1 – 16 h) with the value before laser processing (0 h). The fluorescence intensity shift of the Alexa Fluor 488-BSA gradient was evaluated by subtracting the fluorescence intensity value at the micro-hole distance of 45 – 450 μm to the value at 0 μm.

### Silanization of the thin-glass sheet surface

Silanization aims to change the charge of the thin-glass sheet from negative to positive with organosilane molecules. This process helps the attachment of the cells to the thin-glass sheet. Firstly, the microfluidic device was treated by plasma (Plasma Cleaner CY-P2L-B) at 65 W for 2 min. The surface of the thin-glass sheet was silanized by coating with 2% 3-(2-aminoethylamino)propyldimethoxymethylsilane (Shin-Etsu Chemical Co., Japan) dissolved in pure water for 30 min. The organosilane solution was removed. The coated surface was then baked (As ONE SONW-300) at 80 °C for 1 h to enhance the covalent bonding of the silanol groups. The surface was washed by pure water and 70% ethanol. Finally, the thin-glass sheet embedded in the microfluidic device was ready to be used as culture substrate.

### Cell culture

The silanized thin-glass sheet was coated with poly-D-lysine (PDL) (100 μg/mL) (Gibco, USA) and laminin (50 μg/mL) (Fujifilm Wako Pure Chemical Industries, Ltd, Japan). Hippocampal neurons isolated from embryonic day 16 (E16) mice were provided by the Division of Biological Science, Nara Institute of Science and Technology, Japan. The neurons were seeded on the laminin-coated thin-glass sheet with a cell density of 2.0 × 10^4^ cells/cm^2^. The neurons were cultured in a plate out medium containing Neurobasal (Gibco), 10% fetal bovine serum (FBS) (Gibco), and penicillin (100 units/mL) – streptomycin (100 μg/mL) (Gibco). After 3 h, the medium was changed into a complete medium containing Neurobasal, 2% B-27 supplement (50×) (Gibco), penicillin (100 units/mL) – streptomycin (100 μg/mL). The neurons were cultured in the incubator (37 °C, 5% CO2) for 1 DIV.

### Femtosecond laser processing and time-lapse imaging

Before the manipulation, the netrin-1 (800 ng/mL) was introduced to the channel manually using a micropipette via the inlet. The device containing cultured neurons and netrin-1 was placed under a confocal microscope (Olympus IX71FVSF-2) and maintained in an environmental chamber (Olympus MIIBC; 37 °C, 5% CO2). The fs laser (Spectra-Physics, Solstice-Ref-MT5W, 800 nm, 130 fs) was irradiated through an objective lens (×10, Olympus Plan N, 0.25 NA) on the thin-glass sheet. The frequency and power of 1 kHz and 0.7 μJ/pulse, respectively, were used for fabricating the micro-holes on the thin-glass sheet (Video supplement 1). The response of neurons to the netrin-1 gradient was observed with time-lapse imaging every 10 min using a confocal microscope equipped with a CCD camera (Teli CS3330B), a 10× objective lens, and FLUOVIEW software (Figure 2d).

### Quantification of neuron growth and statistical analysis

The time-lapse pictures of axon elongation and axon angle were measured using ImageJ software (Version 1.52q). Axon angle was the end-point axonal angles toward the micro-holes. The angular change was the turning of axonal angles toward or away from the micro-holes before and during the exposure of molecular gradients (Figure supplement 1). The fluorescence intensity shift of the molecule over time was demonstrated by a linear model and analyzed by regression statistics. Regression statistics showed the slope indicating the relationship between two variables, *p*-value indicating the significant relationship described by the model, and R^2^ indicating the degree of variability of the dependent variable explained by the model. Independent t-test was performed to compare: 1) axonal angles higher and lower than the average turning angle, and 2) guided axon timing, guided axon length, and axon outgrowth rate dependency to axon-to-micro-hole distance.

## Supporting information

Fabrication of holes

Axon length changes with time 1

Axon length changes with time 2

## Author contributions

Dian Anggraini, Conceptualization, Formal analysis, Investigation, Visualization, Methodology, Writing—original draft, Writing—review and editing; Xun Liu, Formal analysis, Investigation; Kazunori Okano, Investigation, Methodology, Writing—review and editing; Yo Tanaka, Resources; Naoyuki Inagaki, Resources, Writing—review & editing; Ming Li, Writing—review & editing; Yoichiroh Hosokawa, Conceptualization, Writing—review & editing; Sohei Yamada, Conceptualization, Writing—original draft, Writing—review and editing; Yaxiaer Yalikun, Conceptualization, Investigation, Methodology, Writing—original draft, Writing—review and editing

## Data availability

All data are available in the manuscript and supporting files.

## Supporting Files

**Figure supplement 1.**
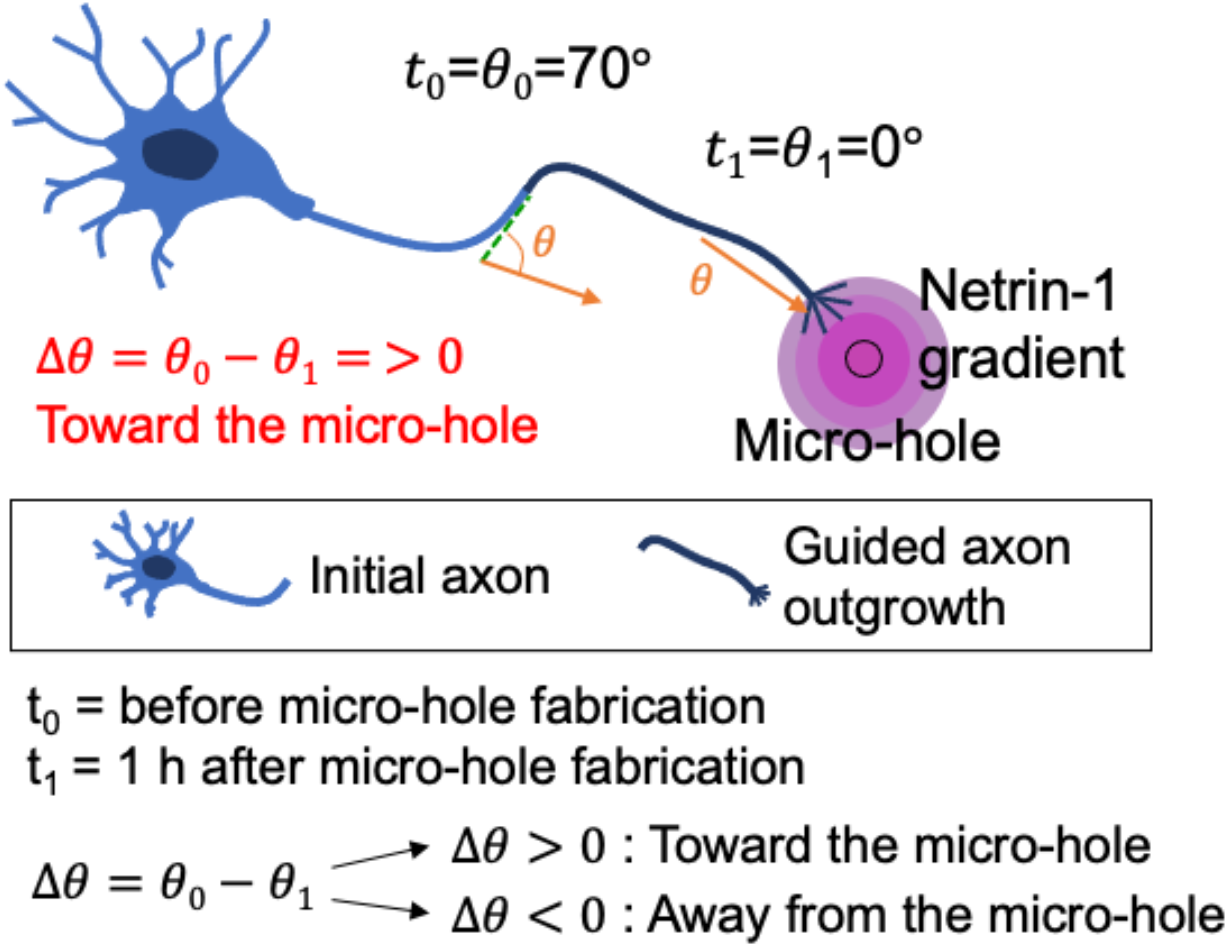
Schematic illustration of the angular change (Δθ) of the initial axon angle (θ_0_) before micro-hole fabrication (t_0_) with axon outgrowth angle (θ_1_) after micro-hole fabrication (t_1_), showing axon outgrowth towards the micro-hole. Angular change (Δ*θ*) is the subtraction of the initial axon angle (*θ*_0_) with axon outgrowth angle (*θ*_1_). If the Δ*θ* >0 and Δ*θ* < 0, it indicates that the axon grows toward and away from the micro-holes, respectively.

**Figure supplement 2.**
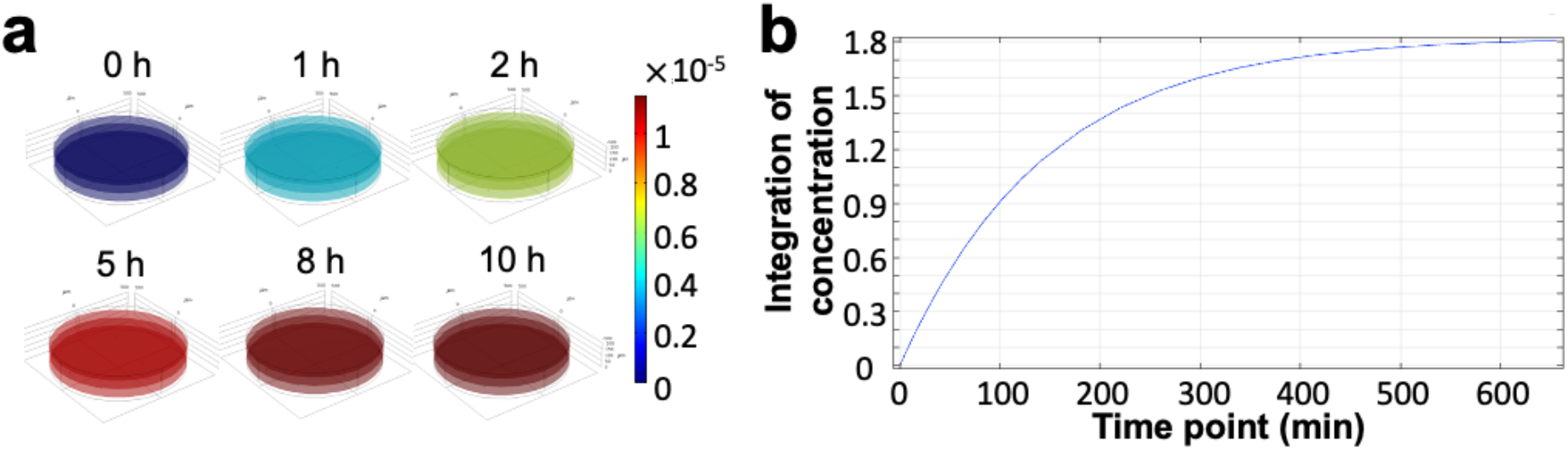
Simulation of molecular gradients. (a) Three-dimensional (3D) diffusion simulation of molecular gradients in the chamber over time. (b) Molecular concentration profile for 10 h from 3D diffusion simulation (Slope = 3.58 × 10^-18^ μm^-1^, R^2^ = 0.86).

**Figure supplement 3.**
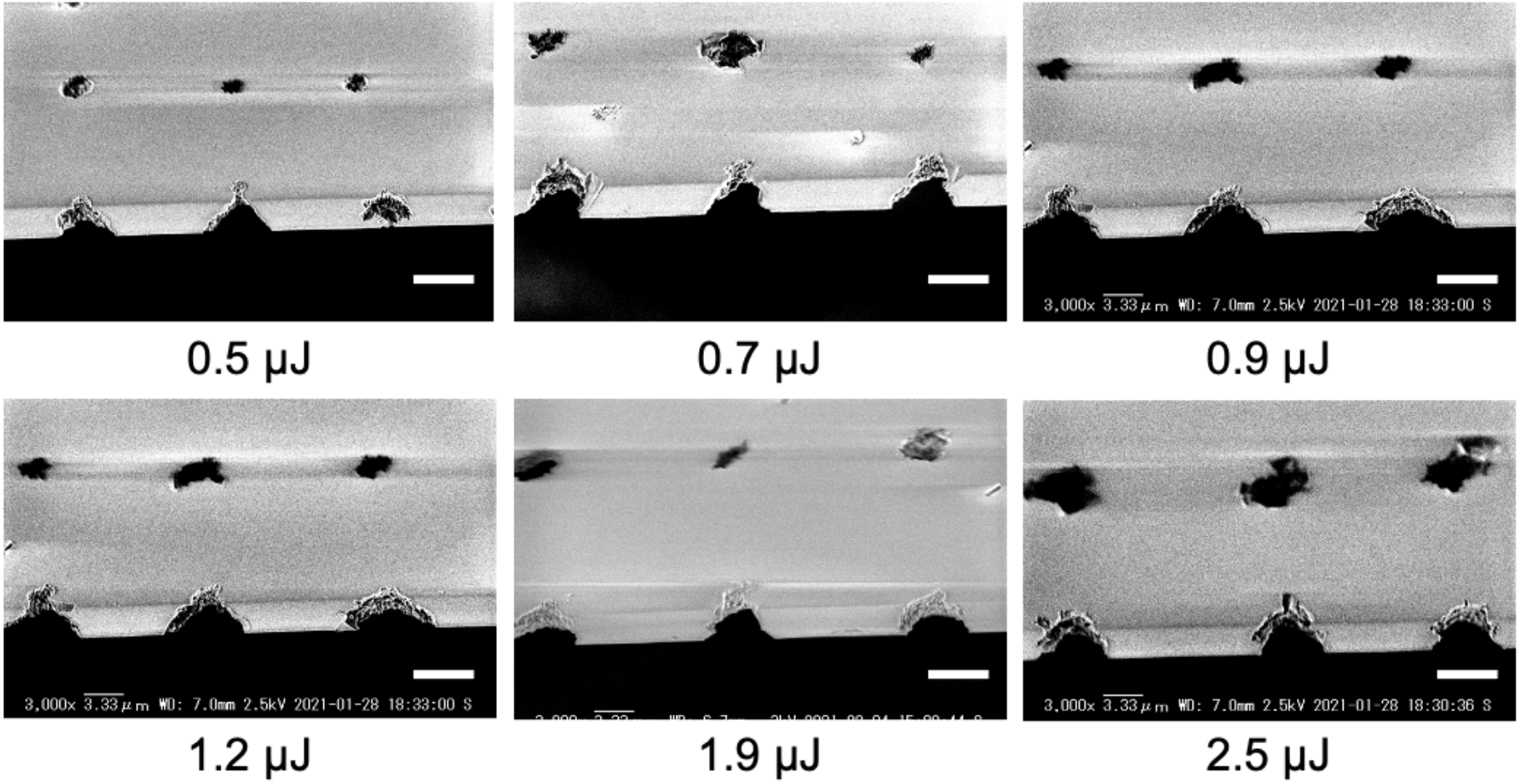
Scanning Electron Microscopy (SEM) images of the 45° view of laser-fabricated micro-holes on the thin-glass sheet fabricated by different pulse energies: 0.5, 0.7, 0.9, 1.2, 1.9, and 2.5 μJ/pulse.

**Figure supplement 4.**
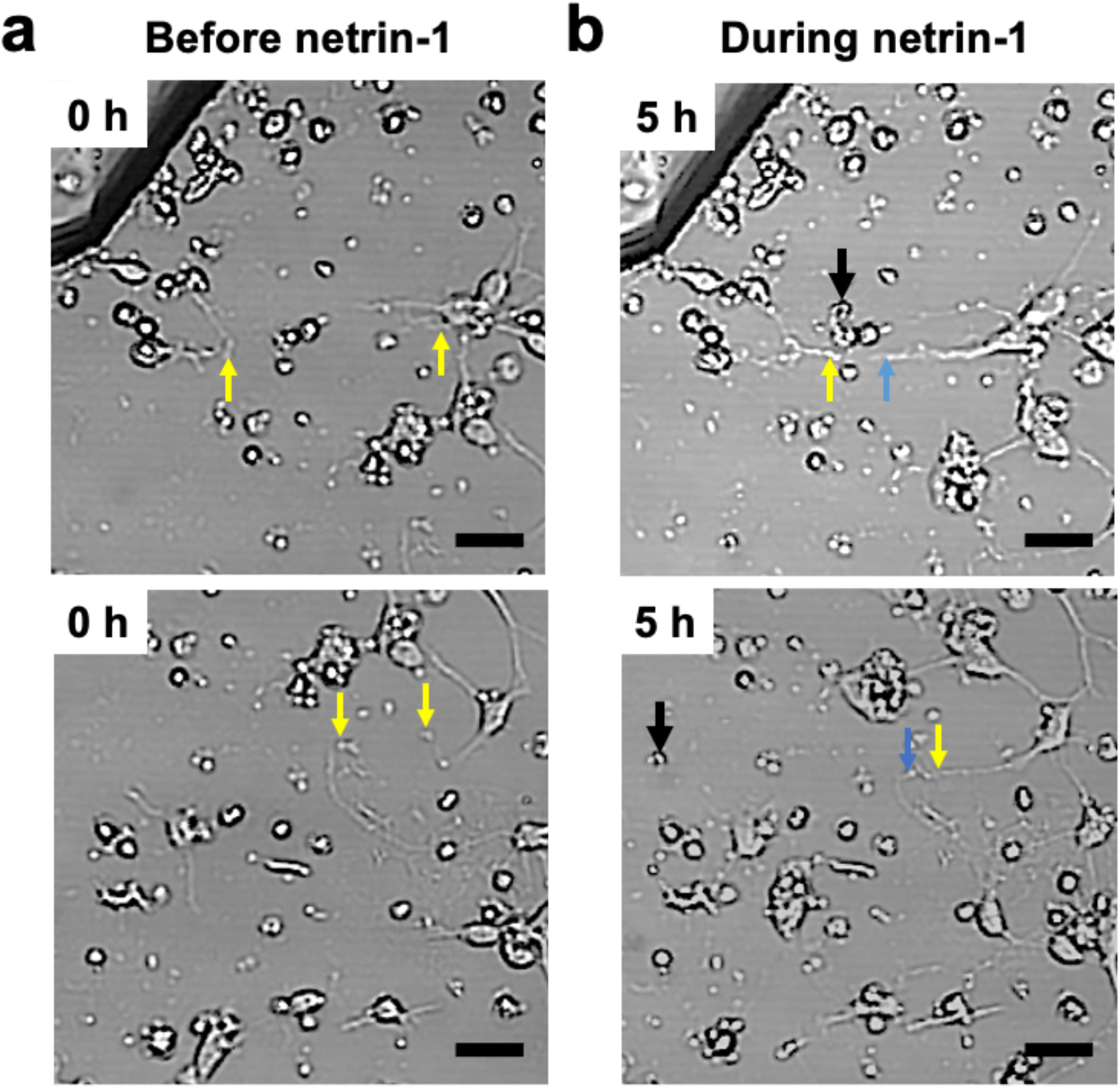
Neuronal culture conditions before and during exposure to netrin-1 released from laser-fabricated micro-holes. (a) Position of initial axons (yellow arrows). (b) The initial axons are guided toward the micro-holes and cross with another guided axon (blue arrow) (up) and preexisting axon (blue arrow) (below). Black arrows are the micro-holes. The scale bars are 50 μm.

**Video supplement 1.** The femtosecond laser pulse of 0.7 μJ/pulse is used for fabricating the micro-hole on the thin-glass sheet during neuronal culture with a short-time laser penetration (2 s in this video) and minimal water bubble generation.

**Video supplement 2.** Axon outgrowth towards the micro-holes for 4 h. The axon is located near the laser-fabricated micro-hole (61.7 μm), showing 61.4 μm axon outgrowth.

**Video supplement 3.** Axon outgrowth towards the micro-holes for 6 h. The axon is located far from the laser-fabricated micro-hole (254 μm), showing 172 μm axon outgrowth.

## References

Alsmadi NZ, Patil LS, Hor EM, Lofti P, Razal JM, Chuong CJ, Wallace GG, Romero-Ortega MI. 2015. Coiled polymeric growth factor gradients for multi-luminal neural chemotaxis. Brain Res 1619:72–83. doi:10.1016/j.brainres.2015.01.055

Baba K, Yoshida W, Toriyama M, Shimada T, Manning CF, Saito M, Kohno K, Trimmer JS, Watanabe R, Inagaki N. 2018. Gradient-reading and mechano-effector machinery for netrin-1-induced axon guidance. Elife 7:1–35. doi:10.7554/eLife.34593

Bhattacharjee N, Folch A. 2017. Large-scale microfluidic gradient arrays reveal axon guidance behaviors in hippocampal neurons. Microsystems Nanoeng 3:17003. doi:10.1038/micronano.2017.3

Bhattacharjee N, Li N, Keenan TM, Folch A. 2010. A neuron-benign microfluidic gradient generator for studying the response of mammalian neurons towards axon guidance factors. Integr Biol 2:669–679. doi:10.1039/c0ib00038h

Blasiak A, Lee GU, Kilinc D. 2015. Neuron Subpopulations with Different Elongation Rates and DCC Dynamics Exhibit Distinct Responses to Isolated Netrin-1 Treatment. ACS Chem Neurosci 6:1578–1590. doi:10.1021/acschemneuro.5b00142

Braisted JE, Catalano SM, Stimac R, Kennedy TE, Tessier-Lavigne M, Shatz CJ, O’Leary DDM. 2000. Netrin-1 promotes thalamic axon growth and is required for proper development of the thalamocortical projection. J Neurosci 20:5792–5801. doi:10.1523/jneurosci.20-15-05792.2000

Carlson AL, Bennett NK, Francis NL, Halikere A, Clarke S, Moore JC, Hart RP, Paradiso K, Wernig M, Kohn J, Pang ZP, Moghe P V. 2016. Generation and transplantation of reprogrammed human neurons in the brain using 3D microtopographic scaffolds. Nat Commun 7:10862. doi:10.1038/ncomms10862

Chen CY, Wo AM, Jong DS. 2012. A microfluidic concentration generator for dose-response assays on ion channel pharmacology. Lab Chip 12:794–801. doi:10.1039/c1lc20548j

Corrigan JD, Selassie AW, Orman JA (Langlois). 2010. The Epidemiology of Traumatic Brain Injury. J Head Trauma Rehabil 25:72–80. doi:10.1097/HTR.0b013e3181ccc8b4

Dinis TM, Elia R, Vidal G, Auffret A, Kaplan DL, Egles C. 2014. Method to form a fiber/growth factor dual-gradient along electrospun silk for nerve regeneration. ACS Appl Mater Interfaces 6:16817–16826. doi:10.1021/am504159j

Dohmen C, Sakowitz OW, Fabricius M, Bosche B, Reithmeier T, Ernestus R-I, Brinker G, Dreier JP, Woitzik J, Strong AJ, Graf R. 2008. Spreading depolarizations occur in human ischemic stroke with high incidence. Ann Neurol 63:720–728. doi:10.1002/ana.21390

Fan Y, Shen F, Frenzel T, Zhu W, Ye J, Liu J, Chen Y, Su H, Young WL, Yang GY. 2010. Endothelial progenitor cell transplantation improves long-term stroke outcome in mice. Ann Neurol 67:488–497. doi:10.1002/ana.21919

Gladkov A, Pigareva Y, Kutyina D, Kolpakov V, Bukatin A, Mukhina I, Kazantsev V, Pimashkin A. 2017. Design of Cultured Neuron Networks in vitro with Predefined Connectivity Using Asymmetric Microfluidic Channels. Sci Rep 7:1–14. doi:10.1038/s41598-017-15506-2

Hawryluk GWJ, Spano S, Chew D, Wang S, Erwin M, Chamankhah M, Forgione N, Fehlings MG. 2014. An examination of the mechanisms by which neural precursors augment recovery following spinal cord injury: A key role for remyelination. Cell Transplant 23:365–380. doi:10.3727/096368912X662408

Hollis ER, Ishiko N, Pessian M, Tolentino K, Lee-Kubli CA, Calcutt NA, Zou Y. 2015. Remodelling of spared proprioceptive circuit involving a small number of neurons supports functional recovery. Nat Commun 6:6079. doi:10.1038/ncomms7079

Homsy A, Laux E, Brossard J, Whitlow HJ, Roccio M, Hahnewald S, Senn P, Mistrík P, Hessler R, Melchionna T, Frick C, Löwenheim H, Müller M, Wank U, Wiesmüller K-H, Keppner H. 2015. Fine control of drug delivery for cochlear implant applications. Hear Balanc Commun 13:153–159. doi:10.3109/21695717.2015.1048082

Hosokawa Y, Iguchi S, Yasukuni R, Hiraki Y, Shukunami C, Masuhara H. 2009. Gene delivery process in a single animal cell after femtosecond laser microinjection. Appl Surf Sci 255:9880–9884. doi:10.1016/j.apsusc.2009.04.111

Hosokawa Y, Ochi H, Iino T, Hiraoka A, Tanaka M. 2011. Photoporation of Biomolecules into Single Cells in Living Vertebrate Embryos Induced by a Femtosecond Laser Amplifier. PLoS One 6:e27677. doi:10.1371/journal.pone.0027677

Ishii N, Wadsworth WG, Stern BD, Culotti JG, Hedgecock EM. 1992. UNC-6, a laminin-related protein, guides cell and pioneer axon migrations in C. elegans. Neuron 9:873–881. doi:10.1016/0896-6273(92)90240-E

Kador KE, Alsehli HS, Zindell AN, Lau LW, Andreopoulos FM, Watson BD, Goldberg JL. 2014. Retinal ganglion cell polarization using immobilized guidance cues on a tissue-engineered scaffold. Acta Biomater 10:4939–4946. doi:10.1016/j.actbio.2014.08.032

Kadoya K, Lu P, Nguyen K, Lee-Kubli C, Kumamaru H, Yao L, Knackert J, Poplawski G, Dulin JN, Strobl H, Takashima Y, Biane J, Conner J, Zhang SC, Tuszynski MH. 2016. Spinal cord reconstitution with homologous neural grafts enables robust corticospinal regeneration. Nat Med 22:479–487. doi:10.1038/nm.4066

Kaji T, Ito S, Miyasaka H, Hosokawa Y, Masuhara H, Shukunami C, Hiraki Y. 2007. Nondestructive micropatterning of living animal cells using focused femtosecond laser-induced impulsive force. Appl Phys Lett 91:023904. doi:10.1063/1.2753103

Kasaai MR, Kacham V, Theberge F, Chin SL. 2003. The interaction of femtosecond and nanosecond laser pulses with the surface of glass. J Non Cryst Solids 319:129–135. doi:10.1016/S0022-3093(02)01909-9

Keino-Masu K, Masu M, Hinck L, Leonardo ED, Chan SS-Y, Culotti JG, Tessier-Lavigne M. 1996. Deleted in Colorectal Cancer (DCC) Encodes a Netrin Receptor. Cell 87:175–185. doi:10.1016/S0092-8674(00)81336-7

Kondziolka D, Wechsler L, Goldstein S, Meltzer C, Thulborn KR, Gebel J, Jannetta P, DeCesare S, Elder EM, McGrogan M, Reitman MA, Bynum L. 2000. Transplantation of cultured human neuronal cells for patients with stroke. Neurology 55:565–569. doi:10.1212/WNL.55.4.565

Kothapalli CR, Van Veen E, De Valence S, Chung S, Zervantonakis IK, Gertler FB, Kamm RD. 2011. A high-throughput microfluidic assay to study neurite response to growth factor gradients. Lab Chip 11:497–507. doi:10.1039/c0lc00240b

Le Harzic R, Huot N, Audouard E, Jonin C, Laporte P, Valette S, Fraczkiewicz A, Fortunier R. 2002. Comparison of heat-affected zones due to nanosecond and femtosecond laser pulses using transmission electronic microscopy. Appl Phys Lett 80:3886–3888. doi:10.1063/1.1481195

Lee BA, Leiby BE, Marino RJ. 2016. Neurological and functional recovery after thoracic spinal cord injury. J Spinal Cord Med 39:67–76. doi:10.1179/2045772314Y.0000000280

Lu P, Wang Y, Graham L, McHale K, Gao M, Wu D, Brock J, Blesch A, Rosenzweig ES, Havton LA, Zheng B, Conner JM, Marsala M, Tuszynski MH. 2012. Long-distance growth and connectivity of neural stem cells after severe spinal cord injury. Cell 150:1264–1273. doi:10.1016/j.cell.2012.08.020

Ma VY, Chan L, Carruthers KJ. 2014. Incidence, Prevalence, Costs, and Impact on Disability of Common Conditions Requiring Rehabilitation in the United States: Stroke, Spinal Cord Injury, Traumatic Brain Injury, Multiple Sclerosis, Osteoarthritis, Rheumatoid Arthritis, Limb Loss, and Back Pa. Arch Phys Med Rehabil 95:986–995. doi:10.1016/j.apmr.2013.10.032

Millet LJ, Stewart ME, Nuzzo RG, Gillette MU. 2010. Guiding neuron development with planar surface gradients of substrate cues deposited using microfluidic devices. Lab Chip 10:1525–1535. doi:10.1039/c001552k

Ming GL, Song HJ, Berninger B, Holt CE, Tessier-Lavigne M, Poo MM. 1997. cAMP-dependent growth cone guidance by netrin-1. Neuron 19:1225–1235. doi:10.1016/S0896-6273(00)80414-6

Moore SW, Biais N, Sheetz MP. 2009. Traction on immobilized netrin-1 is sufficient to reorient axons. Science (80-) 325:166. doi:10.1126/science.1173851

Moore SW, Zhang X, Lynch CD, Sheetz MP. 2012. Netrin-1 Attracts Axons through FAK-Dependent Mechanotransduction. J Neurosci 32:11574–11585. doi:10.1523/JNEUROSCI.0999-12.2012

Nagel AN, Marshak S, Manitt C, Santos RA, Piercy MA, Mortero SD, Shirkey-Son NJ, Cohen-Cory S. 2015. Netrin-1 directs dendritic growth and connectivity of vertebrate central neurons in vivo. Neural Dev 10:14. doi:10.1186/s13064-015-0041-y

Nam SW, Noort D Van, Yang Y, Park S. 2007. A biological sensor platform using a pneumatic-valve controlled microfluidic device containing Tetrahymena pyriformis. Lab Chip 7:638–640. doi:10.1039/b617357h

Nayak TR, Jian L, Phua LC, Ho HK, Ren Y, Pastorin G. 2010. Thin films of functionalized multiwalled carbon nanotubes as suitable scaffold materials for stem cells proliferation and bone formation. ACS Nano 4:7717–7725. doi:10.1021/nn102738c

Okano K, Liu LL, Hosokawa Y, Masuhara H. 2016. In situ dynamic control of neurite growth by femtosecond laser ablation of substrate patternsMHS 2016 - International Symposium on Micro-NanoMechatronics and Human Science, Nagoya, Japan, 30 November 2016. doi:10.1109/MHS.2016.7824186

Okano K, Matsui A, Maezawa Y, Hee PY, Matsubara M, Yamamoto H, Hosokawa Y, Tsubokawa H, Li YK, Kao FJ, Masuhara H. 2013. In situ laser micropatterning of proteins for dynamically arranging living cells. Lab Chip 13:4078–4086. doi:10.1039/c3lc50750e

Okano K, Yu D, Matsui A, Maezawa Y, Hosokawa Y, Kira A, Matsubara M, Liau I, Tsubokawa H, Masuhara H. 2011. Induction of Cell-Cell Connections by Using in situ Laser Lithography on a Perfluoroalkyl-Coated Cultivation Platform. ChemBioChem 12:795–801. doi:10.1002/cbic.201000497

Oyama K, Zeeb V, Kawamura Y, Arai T, Gotoh M, Itoh H, Itabashi T, Suzuki M, Ishiwata S. 2015. Triggering of high-speed neurite outgrowth using an optical microheater. Sci Rep 5:16611. doi:10.1038/srep16611

Pearse DD, Pereira FC, Marcillo AE, Bates ML, Berrocal YA, Filbin MT, Bunge MB. 2004. cAMP and Schwann cells promote axonal growth and functional recovery after spinal cord injury. Nat Med 10:610–616. doi:10.1038/nm1056

Pedersen PM, Stig Jørgensen H, Nakayama H, Raaschou HO, Olsen TS. 1995. Aphasia in acute stroke: Incidence, determinants, and recovery. Ann Neurol 38:659–666. doi:10.1002/ana.410380416

Prabhakaran MP, Venugopal JR, Chyan T Ter, Hai LB, Chan CK, Lim AY, Ramakrishna S. 2008. Electrospun biocomposite nanofibrous scaffolds for neural tissue engineering. Tissue Eng - Part A 14:1787–1797. doi:10.1089/ten.tea.2007.0393

Riess P, Zhang C, Saatman KE, Laurer HL, Longhi LG, Raghupathi R, Lenzlinger PM, Lifshitz J, Boockvar J, Neugebauer E, Snyder EY, McIntosh TK. 2002. Transplanted Neural Stem Cells Survive, Differentiate, and Improve Neurological Motor Function after Experimental Traumatic Brain Injury. Neurosurgery 51:1043–1054. doi:10.1097/00006123-200210000-00035

Rukmana TI, Moran G, Méallet-Renault R, Ohtani M, Demura T, Yasukuni R, Hosokawa Y. 2019. Enzyme-Assisted Photoinjection of Megadalton Molecules into Intact Plant Cells Using Femtosecond Laser Amplifier. Sci Rep 9:1–9. doi:10.1038/s41598-019-54124-y

Serafini T, Kennedy TE, Gaiko MJ, Mirzayan C, Jessell TM, Tessier-Lavigne M. 1994. The netrins define a family of axon outgrowth-promoting proteins homologous to C. elegans UNC-6. Cell 78:409–424. doi:https://doi.org/10.1016/0092-8674(94)90420-0

Shear DA, Tate MC, Archer DR, Hoffman SW, Hulce VD, Laplaca MC, Stein DG. 2004. Neural progenitor cell transplants promote long-term functional recovery after traumatic brain injury. Brain Res 1026:11–22. doi:10.1016/j.brainres.2004.07.087

Shi L, Kuhnell D, Borra VJ, Langevin SM, Nakamura T, Esfandiari L. 2019. Rapid and label-free isolation of small extracellular vesicles from biofluids utilizing a novel insulator based dielectrophoretic device. Lab Chip 19:3726–3734. doi:10.1039/c9lc00902g

Shin JC, Kim KN, Yoo J, Kim I-S, Yun S, Lee H, Jung K, Hwang K, Kim M, Lee I-S, Shin JE, Park KI. 2015. Clinical Trial of Human Fetal Brain-Derived Neural Stem/Progenitor Cell Transplantation in Patients with Traumatic Cervical Spinal Cord Injury. Neural Plast 2015:1–22. doi:10.1155/2015/630932

Tang T, Yuan Y, Yalikun Y, Hosokawa Y, Li M, Tanaka Y. 2021. Glass based micro total analysis systems: Materials, fabrication methods, and applications. Sensors Actuators, B Chem 339:129859. doi:10.1016/j.snb.2021.129859

Tao S, Young C, Redenti S, Zhang Y, Klassen H, Desai T, Young MJ. 2007. Survival, migration and differentiation of retinal progenitor cells transplanted on micro-machined poly(methyl methacrylate) scaffolds to the subretinal space. Lab Chip 7:695–701. doi:10.1039/b618583e

Tao SL, Desai TA. 2007. Aligned arrays of biodegradable poly(ε-caprolactone) nanowires and nanofibers by template synthesis. Nano Lett 7:1463–1468. doi:10.1021/nl0700346

Taylor AM, Dieterich DC, Ito HT, Kim SA, Schuman EM. 2010. Microfluidic Local Perfusion Chambers for the Visualization and Manipulation of Synapses. Neuron 66:57–68. doi:10.1016/j.neuron.2010.03.022

Taylor AM, Menon S, Gupton SL. 2015. Passive microfluidic chamber for long-term imaging of axon guidance in response to soluble gradients. Lab Chip 15:2781–2789. doi:10.1039/c5lc00503e

Vaysse L, Beduer A, Sol JC, Vieu C, Loubinoux I. 2015. Micropatterned bioimplant with guided neuronal cells to promote tissue reconstruction and improve functional recovery after primary motor cortex insult. Biomaterials 58:46–53. doi:10.1016/j.biomaterials.2015.04.019

Vroemen M, Aigner L, Winkler J, Weidner N. 2003. Adult neural progenitor cell grafts survive after acute spinal cord injury and integrate along axonal pathways. Eur J Neurosci 18:743–751. doi:10.1046/j.1460-9568.2003.02804.x

Xiao RR, Zeng WJ, Li YT, Zou W, Wang L, Pei XF, Xie M, Huang WH. 2013. Simultaneous generation of gradients with gradually changed slope in a microfluidic device for quantifying axon response. Anal Chem 85:7842–7850. doi:10.1021/ac4022055

Xu Z, Fang P, Xu B, Lu Y, Xiong J, Gao F, Wang X, Fan J, Shi P. 2018. High-throughput three-dimensional chemotactic assays reveal steepness-dependent complexity in neuronal sensation to molecular gradients. Nat Commun 9:4745. doi:10.1038/s41467-018-07186-x

Yalikun Y, Hosokawa Y, Iino T, Tanaka Y. 2016. An all-glass 12 μm ultra-thin and flexible micro-fluidic chip fabricated by femtosecond laser processing. Lab Chip 16:2427–2433. doi:10.1039/c6lc00132g

Yalikun Y, Tanaka N, Hosokawa Y, Iino T, Tanaka Y. 2017. Embryonic body culturing in an all-glass microfluidic device with laser-processed 4 μm thick ultra-thin glass sheet filter. Biomed Microdevices 19:1–10. doi:10.1007/s10544-017-0227-7

Yalikun Y, Tanaka Y. 2017. Ultra-thin glass sheet integrated transparent diaphragm pressure transducer. Sensors Actuators, A Phys 263:102–112. doi:10.1016/j.sna.2017.05.047

Yamamoto H, Okano K, Demura T, Hosokawa Y, Masuhara H, Tanii T, Nakamura S. 2011. In-situ guidance of individual neuronal processes by wet femtosecond-laser processing of self-assembled monolayers. Appl Phys Lett 99:163701. doi:10.1063/1.3651291

Yang TC, Chuang JH, Buddhakosai W, Wu WJ, Lee CJ, Chen WS, Yang YP, Li MC, Peng CH, Chen SJ. 2017. Elongation of Axon Extension for Human iPSC-Derived Retinal Ganglion Cells by a Nano-Imprinted Scaffold. Int J Mol Sci 18:2013. doi:10.3390/ijms18092013

Yokota K, Kobayakawa K, Kubota K, Miyawaki A, Okano H, Ohkawa Y, Iwamoto Y, Okada S. 2015. Engrafted Neural Stem/Progenitor Cells Promote Functional Recovery through Synapse Reorganization with Spared Host Neurons after Spinal Cord Injury. Stem Cell Reports 5:264–277. doi:10.1016/j.stemcr.2015.06.004

Yuan Y, Yalikun Y, Amaya S, Aishan Y, Shen Y, Tanaka Y. 2021. Fabrication of ultra-thin glass sheet by weight-controlled load-assisted precise thermal stretching. Sensors Actuators, A Phys 321:112604. doi:10.1016/j.sna.2021.112604

Zhang S, Wang XJ, Li WS, Xu XL, Hu JB, Kang XQ, Qi J, Ying XY, You J, Du YZ. 2018. Polycaprolactone/polysialic acid hybrid, multifunctional nanofiber scaffolds for treatment of spinal cord injury. Acta Biomater 77:15–27. doi:10.1016/j.actbio.2018.06.038

